# Exploring population responses to environmental change when there’s never enough data; a factor analytic approach

**DOI:** 10.1101/184036

**Authors:** Bethan J. Hindle, Mark Rees, Andy W. Sheppard, Pedro F. Quintana-Ascencio, Eric S. Menges, Dylan Z. Childs

## Abstract

1. Temporal variability in the environment drives variation in individuals’ vital rates, with consequences for population dynamics and life history evolution. Integral projection models (IPMs) are data-driven models widely used to study population dynamics and life history evolution of structured populations in temporally variable environments. However, many data sets have insufficient temporal replication for the environmental drivers of vital rates to be identified with confidence, limiting their use for evaluating population level responses to environmental change.
2. Parameter selection, where the kernel is constructed at each time step by randomly selecting the time-varying parameters from their joint probability distribution, is one approach to including stochasticity in IPMs. We consider a factor analytic (FA) approach for modelling the covariance matrix of time-varying parameters, whereby latent variable(s) describe the covariance among vital rate parameters. This decreases the number of parameters to estimate and, where the covariance is positive, the latent variable can be interpreted as a measure of environmental quality. We demonstrate this using simulation studies and two case studies.
3. The simulation studies suggest the FA approach provides similarly accurate estimates of stochastic population growth rate to estimating an unstructured covariance matrix. We demonstrate how the latent parameter can be perturbed to show how selection on reproductive delays in the monocarp *Carduus nutans* changes under different environmental conditions. We develop a demographic model of the fire dependent herb *Eryngium cuneifolium* to show how a causal indicator (i.e. a driver of the changes in the environmental quality) can be incorporated with the addition of a single parameter. Using perturbation analyses we determine optimal management strategies for this species.
4. This approach estimates fewer parameters than previous approaches and allows novel eco-evolutionary insights. Predictions on population dynamics and life history evolution under different environmental conditions can be made without necessarily identifying causal factors. Environmental drivers can be incorporated with relatively few parameters, allowing for predictions on how populations will be affected by changes to these drivers.

## Introduction

Environmental variation causes individuals’ vital rates to vary, affecting population dynamics and life history evolution (Benton & Grant 1996; Boyce *et al*. 2006). Interest in understanding the ecological consequences of environmental variation has increased rapidly as a consequence of global climate change (Stenseth *et al*. 2002; Evans 2012). As experimental approaches to determining how natural populations are affected by environmental variation are frequently impractical, structured demographic models are often used to understand the population level effects of environmental change (Coulson 2012). Environmental effects on vital rates can be complex, with nonlinear effects, multiple interacting drivers, indirect effects and correlations between the drivers (Darling & Cote 2008; Parmesan *et al*. 2013; Ehrlen *et al*. 2016). These challenges, and the relatively short length of many demographic data sets (Salguero-Gomez *et al*. 2015; Salguero-Gomez *et al*. 2016), mean it is often difficult to identify explicit environmental drivers of vital rates. This restricts the ability of models to predict how populations will respond environmental change (Crone *et al*. 2013).

Environmental variation can drive covariation amongst vital rates (Doak *et al*. 2005; Tomimatsu & Ohara 2010). All else equal, failing to account for this covariation will bias model outputs (Fieberg & Ellner 2001; Metcalf *et al*. 2015). Positive covariance among vital rates, occurring when multiple vital rates are affected by the same environmental drivers (Jongejans *et al*. 2010), increases the variance of the stochastic population growth rate. Negative covariance can also occur as a result of tradeoffs between rates or from opposing effects of environmental variables on different rates (Jongejans & De Kroon 2005; Knops, Koenig & Carmen 2007). However, in plants covariation is predominantly positive (Jongejans *et al*. 2010), and positive covariance appears widespread among other taxa including mammals (e.g. Rotella *et al*. 2012) and birds (e.g. Jenkins, Watson & Miller 1963; Nur & Sydeman 1999).

Stochastic demographic models, such as matrix population models (MPMs; see Caswell 2001) and integral projection models (IPMs; see Ellner, Childs & Rees 2016), are widely used to study population dynamics under temporally variable environments (e.g. Inchausti & Weimerskirch 2001; Vindenes *et al*. 2014). IPMs are usually parameterised by estimating state-fate relationships. Stochastic models allow these relationships to vary temporally (or spatially), using one of two main methods (Metcalf *et al*. 2015). Under a kernel selection approach, a projection kernel is estimated for each year and these are resampled (Rees *et al*. 2006; Williams *et al*. 2015) to preserve the covariance amongst the vital rates. Using a parameter selection approach, a unique kernel is constructed at each time step by randomly selecting the time-varying parameters from their joint probability distribution (Morris & Doak 2002; Vindenes *et al*. 2014). A potential limitation of the parameter selection approach is that an unstructured covariance matrix must be estimated for the set of time varying parameters, often from relatively few temporal replicates.

An alternative to estimating an unstructured covariance matrix is to use a structured model for the temporal parameters (co)variances. Hierarchical factor analysis (FA), whereby one or more latent variables are introduced to capture the covariance among vital rate parameters, is a promising candidate. The latent variable(s) represent the underlying causes of covariation among observed variables, allowing complex multivariate relationships to be described in a simple way. Moreover, these models effectively capture hypotheses about causal variables that cannot be directly measured (Grace & Bollen 2008; Grace *et al*. 2010). The FA approach can be also be extended to include the underlying drivers of variation in the latent variables as causal indicators, which allows covariances to be partitioned into explained and unexplained sources of variation. However, despite the broad use of FA approaches in ecological research (e.g. Zuur *et al*. 2003; Thorson *et al*. 2015; Ohlberger, Scheuerell & Schindler 2016) they are rarely used to parameterise demographic models.

This approach has two potential advantages. First, fewer parameters need to be estimated relative to an unstructured covariance matrix. Second, a small number of latent variables (often just one) may account for the covariation among the vital rates. When this covariance is positive, the latent variable(s) can be interpreted as axes of environmental quality or suitability, where positive values of a single latent variable correspond to environments in which survival, growth and reproduction are all higher than average. The latent term(s) then represent a target for further analysis. For example, perturbing the latent parameter allows predictions to be made on the effects of environmental change on population dynamics or life history selection. Where the degree of temporal replication in the data is insufficient for causal environmental drivers to be identified this may represent the best alternative for exploring how changes in the stochastic part of the environment affect such processes. This method is not dissimilar to the use of broad scale climate indices, such as the North Atlantic Oscillation, as proxies for local environmental conditions (Ottersen *et al*. 2001). Such indices do not directly influence the vital rates, but as they provide an index of the overall climate conditions, incorporating multiple local climate variables, they are often better predictors of the vital rates than local climate variables (Post & Stenseth 1999; Stenseth & Mysterud 2005).

We conduct simulation studies to compare the accuracy of the FA approach to a standard parameter selection approach, with different numbers of temporally varying parameters. We then apply the approach in two case studies. We construct a demographic model of the monocarpic perennial *Carduus nutans*, and show how the latent parameter can be perturbed to make predictions about optimal life history strategies under changing environments. We explore how selection for strategies to delay reproduction differs as the mean and variance of environmental quality changes. Finally, we develop a demographic model of the rare herb *Eryngium cuneifolium* to show how a known environmental driver (time since fire) can be included as a causal indicator, i.e. an observed variable that influences the latent variable. We use perturbation analyses to determine the optimal fire return interval (FRI) for managing this species.

### Simulation study: comparing factor analytic and unstructured approaches

We compared the accuracy of population growth estimates from the FA approach to those derived using an unstructured covariance matrix. We considered two scenarios: a relatively simple life history with four temporally variable vital rates (the ‘simple model’), typical of many published IPMs, and a two-stage (juvenile and adult) life history with a total of seven temporally variable vital rates (the ‘complex model’). Demographic rate functions in both settings were parameterised using data from a long-term study of the St Kilda Soay sheep (Clutton-Brock & Pemberton 2004). These were used to construct a pair of density independent individual-based models (IBMs; Appendix A1), from which simulated data sets could be generated. Only the correlation coefficients for the temporally varying parameters were allowed to vary in each simulation, such that on each occasion, a correlation matrix was drawn at random from a uniform distribution over the space of positive definite matrices (using rcorrmatrix from the clusterGeneration package in R Qiu & Joe 2015).

One hundred simulated data sets of 8,000 years were generated from each of the two IBMs. A range of realistic data set lengths were sampled: 12, 25 and 50 years (see Appendix A1 for details). To account for the covariance among vital rates, multivariate demographic models were then parameterised using an unstructured covariance matrix (UCM approach) and a latent variable (FA approach) parameterisation (Fig. 1). In the simple model, 10 parameters (4 variance and 6 covariances) account for the temporal variation using the UCM approach, whilst the FA approach estimates 8 parameters. In the complex model 28 parameters are required for the UCM approach and 14 for the FA approach. The demographic models were fitted using Bayesian methods, implemented in JAGS (Plummer 2003) and run using the runjags package (Denwood in review) in R (R Core Team 2016).

**Figure 1:**
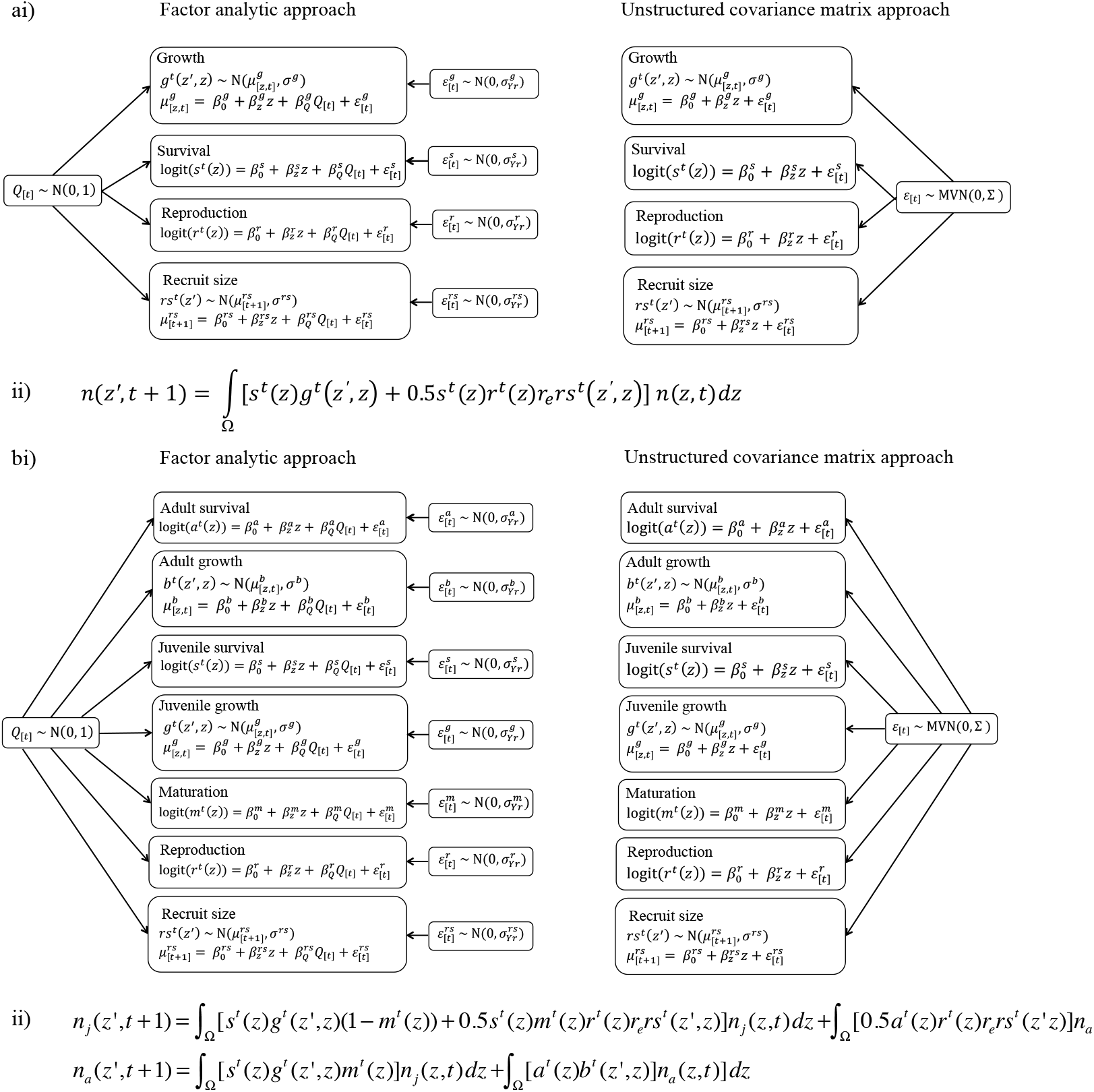
Structure of the i) vital rate models and ii) IPMs for the a) simple and b) complex life history simulation models. In the FA approach factor-loading terms (*β_Q_*) allowed the direction and magnitude of the latent parameter (*Q*) to differ among the vital rates. Submodel specific year effects (*ε*) accounted for any additional variation among years. In the UCM approach a fully unstructured covariance matrix (Σ) was estimated by sampling the year effects (*ε*) from a multivariate normal distribution. *β*_0_ parameters are the intercepts and *β_z_* are slopes with respect to size. aii) is the structure of the IPM for the simple life history model, where *n(z, t)* is the size distribution of individuals at time *t, s^t^(z)g^t^(z',z)* is a survival growth kernel and 0.5*s^t^(z)r^t^(z)r_e_rs^t^(z',z)* gives the size distribution of new recruits. The superscript *t* denotes stochastic terms. bii) is the structure of the IPM for the complex model. The size distribution of juveniles at time t, *n_j_(z, t)*, is given by the survival and growth of juveniles at *t* - 1 that do not mature that year, *s^t^(z)g^t^(z',z)(1-m^t^(z)*, and the reproduction of adults, 0.5*a^t^(z)r^t^(z)r_e_rs^t^(z',z)*, and juveniles that have matured that year, 0.5*s^t^(z)m^t^(z)r^t^(z)r^t^(z)r_e_c^t^(z',z)*. The size distribution of adults at time *t, n_a_(z, t)*, is given by the survival-growth function of maturing juveniles, *s^t^(z)g^t^(z',z)m^t^(z)*, and the survival growth function of adults, *a^t^(z)b^t^(z', z)*. In both a) and b) the functions in the IPMs correspond to the functions in i), with the exception of lamb recruitment (*r_e_*), which is assumed not to vary with time (Appendix A1).

IPMs were constructed from each set of posterior samples (Fig. 1; Appendix A1). The stochastic population growth rate was estimated after excluding the first 2,000 years of a 10,000 year simulation. This was repeated with 1,000 samples from the posterior. The true stochastic population growth rate was estimated using an IPM parameterised with the true parameter values used in the IBM.

The results of the simulation study are summarised in Fig. 2. The UCM approach led to marginally less diffuse estimates of stochastic population growth rate than the FA approach. This was true for both the simple (Fig. 2a) and complex (Fig. 2b) models. However, even with 12 years of temporal replication the differences between the performance of the two methods was small, and with 25 years of replication both methods performed well.

**Figure 2:**
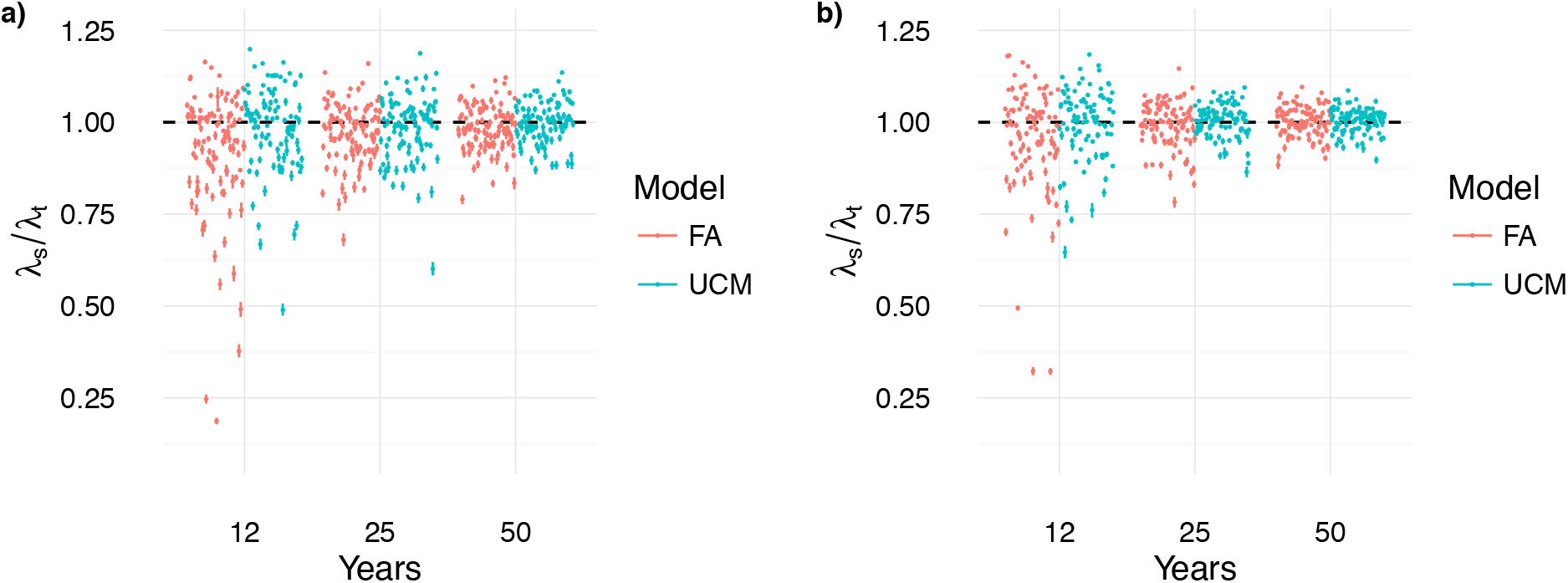
Ratio between the true (*λ_t_*) and estimated (*λ_s_*) stochastic population growth rates for the factor analytic (FA) and unstructured covariance matrix (UCM) approaches for the a) simple and b) complex models. Points are the median and lines show the 0.025 and 0.975 quantiles across 1000 samples from the posteriors for each simulation. The dashed line is at one, where the estimated growth rate equals the true growth rate.

### Case study 1: The effect of environmental quality on reproductive delays in *Carduus nutans*

#### Background and methods

*Carduus nutans* is a monocarpic thistle with a persistent seedbank and short-lived rosettes (Popay & Medd 1990; Wardle, Nicholson & Rahman 1992). We use a FA model to explore how environmental change may affect selection for reproductive delays in this species. Reproductive delays can act as a form of diversified bet hedging, spreading a cohort across multiple years and therefore decreasing the effect of a bad year on the cohort as a whole (Cohen 1966; Tuljapurkar 1990; Rees *et al*. 2006; Childs, Metcalf & Rees 2010). In monocarpic perennial plants, reproduction may be deferred pre-establishment, through a seedbank, or post-establishment, through a delay in flowering (Childs *et al*. 2004; Rees *et al*. 2006). Post-establishment delays have the additional benefit of higher fecundity as individuals may grow larger, producing more seeds (Rees *et al*. 2006).

We define the fittest strategy to be the evolutionary stable strategy (ESS). The predicted ESS for the study population is substantial seed dormancy and the majority of plants to flower in their first year, with a flowering probability of ~0.75 for an average sized individual (Rees *et al*. 2006). Using our framework we predict how changes to the average or variability of the environment affect the ESS germination and flowering strategy. We re-parameterised the IPM of Rees et al (2006; Fig. 3a, Appendix A2). The model is structured by the natural logarithm of rosette area (*z*), a measure of plant size that predicts individual performance. Four stochastic vital rate functions, with temporally variable intercepts, were estimated; survival, growth, recruitment and recruit size (Fig. 3).

**Figure 3:**
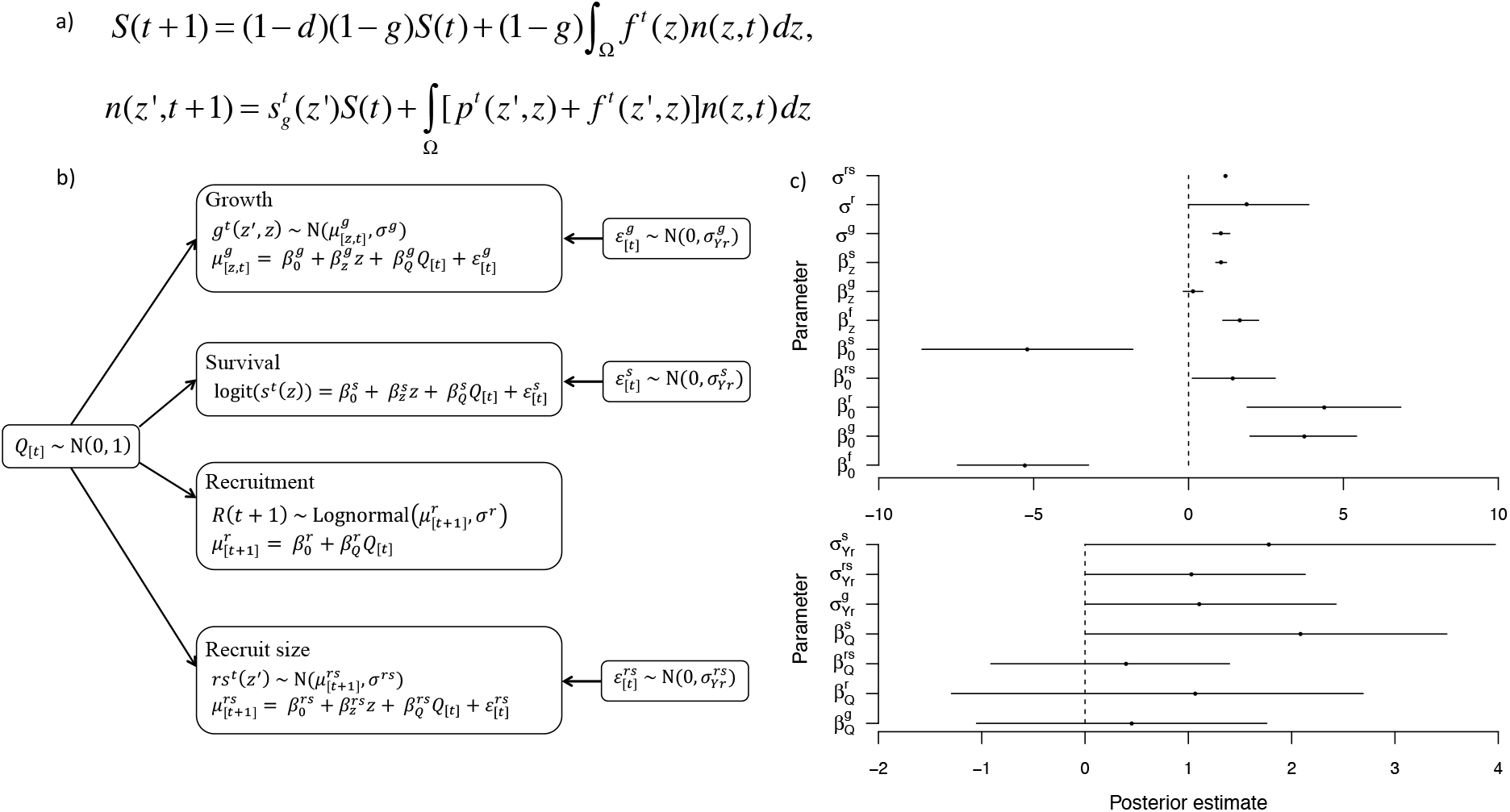
a) *Carduus* IPM kernel (see Appendix A2), where () is the number of seeds in the seedbank and *n(z', t)dz* is the size distribution of rosettes in year *t*. The first term in the seedbank equation equates to seeds present in the seedbank at time *t* that have not died (with probability of seed mortality *d*) or germinated (with probability of germination *g*). The second term is seeds produced by rosettes, where *f^t^(z)* is a seed production function. The rosette equation can be split into three terms, describing seeds germinating from the seedbank 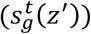, rosette survival and growth (*p^t^(z',z)*) and seeds produced that year that germinate immediately (*f^t^(z', z))*. The superscript denotes stochastic terms. b) Structure of the stochastic vital rates model, including equations for each submodel, and c) posterior distributions for (i) fixed parameters and (ii) parameters for incorporating temporal variation. The subscripts [*t*] indicate stochastic terms. *β*_0_ parameters are the intercepts, *β_z_* and *β_Q_* parameters are slopes with respect to size (*z*) and the latent parameter (*Q*) respectively and ε are submodel specific year effects. 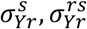 and 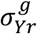 are the standard deviations of the submodel specific year effects for survival, recruit size and growth respectively. Recruitment refers to the number of seedlings at the annual census; as this is a single number each year an additional year effect is not included here. c) shows the mean (points) and 95% credible interval (horizontal lines) for each parameter. Vertical dashed lines are at 0.

The vital rate parameters (Fig. 3b) were estimated using MCMC sampling in JAGS through runjags (Denwood in review; see example JAGs code in supplementary material). The prior distributions were weakly informative (i.e. within biologically reasonable ranges) to improve mixing (Appendix A2; see Appendix A3 for comparison with more informative priors). The vital rates were integrated into the IPM (Fig. 3) using the posterior means as parameter estimates. At each year in the simulation the latent parameter (*Q*) was sampled from a normal distribution with a mean of zero and a standard deviation of one. The submodel specific year effects (*ε*) were drawn from normal distributions with means of zero and the standard deviations (*σ_Yr_*) estimated in the vital rates model.

Posterior checks suggested the latent parameter (*Q*) accounted for the covariation among the vital rates (Fig. S1). The 95% credible intervals of many parameters were relatively wide (Fig. 3c), as a result of the short temporal extent (eight years) of this data set. The positive covariance among the vital rates (Fig. 3 & S2) means the latent parameter (*Q*) can be assumed to be a measure of environmental quality. The highest levels of temporal variation were in survival and recruitment (Fig. S2). The joint flowering intercept and germination probability ESS were predicted using numerical invasion analysis (Childs *et al*. 2004) and were similar to those produced using a fixed effects, kernel selection approach (Appendix A4; Rees *et al*. 2006).

#### Perturbation analyses

A prospective sensitivity analysis was used to determine how selection on delayed flowering and germination may change as the mean and variance of environmental quality (*Q*) changes. The mean and standard deviations of *Q* were varied on a fixed grid and the ESS were predicted at each value. This was repeated for a range of seed mortalities (d=0.01, 0.1, 0.2…0.9, 0.99; Rees *et al*. 2006).

As the quality of the environment deteriorates there is selection for earlier flowering and reduced germination, whilst improving the quality of the environment leads to the opposite response, i.e. selection for a perennial life history dominates in higher quality environments (Figs 4 & S3). In lower quality environments selection acts on the germination probability, delaying reproduction pre-establishment by increasing the chance of seeds entering the seedbank. The estimated survival probability increases from 0.04 to 0.73 with an increase in *Q* from 0 to 2 for a rosette of log size 1.95 (mode of the study population). With a mean *Q* of 2 there is an advantage in delaying reproduction, as the risk of mortality is relatively small and larger plants can produce more seeds; here, selection acts on the flowering size, increasing the size at which plants reproduce. The ESS threshold flowering size, on a log scale, doubles from 3.36 to 7.10 with an increase in mean *Q* from 0 to 2, resulting in a 9-fold increase in the estimated number of seeds produced. Increasing levels of environmental variability generally caused selection for earlier flowering and a lower germination probability (Fig. 4).

**Figure 4:**
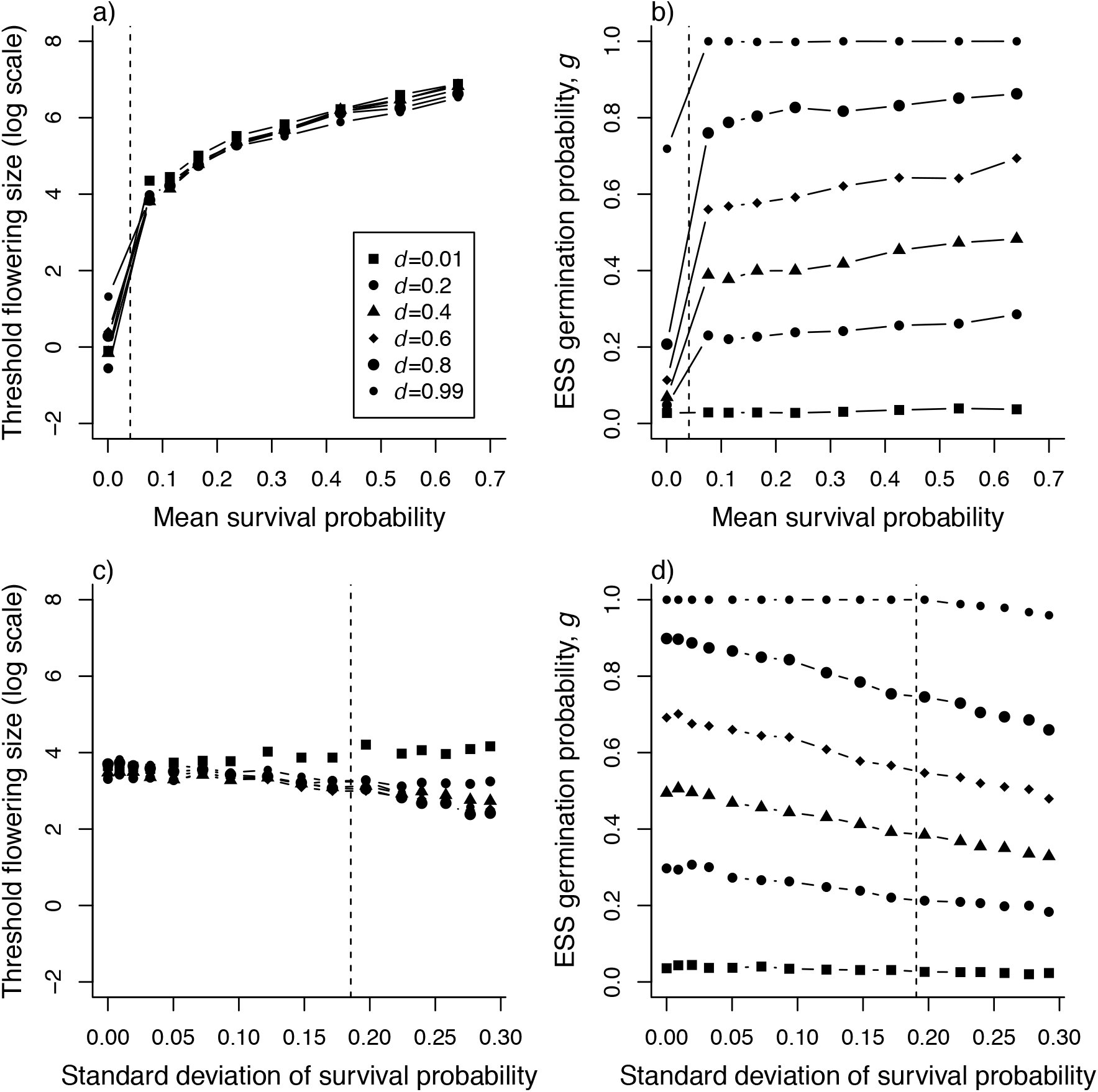
Effect of changing the mean (a & b) and variability (c & d) of the environmental quality (*Q*) on the joint ESS flowering intercept and germination strategies at different levels of seed mortality (*d*) in *Carduus*. Changes to the environmental quality here are shown through the impact on survival of an averaged sized individual (log size 1.95, which is the mode of the study population). Threshold flowering size is calculated as 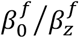 (Childs *et al*. 2003). Dashed lines show the mean survival probability (a and b) and standard deviation of the survival probability (c and d) for the average sized individual in the environment experienced by the study population.

### Case study 2: Incorporating an environmental driver: the effect of fire on the demography of *Eryngium cuneifolium*

#### Background and methods

*Eryngium* is a fire-adapted perennial herb with a persistent seedbank (Menges & Kimmich 1996; Menges & Quintana-Ascencio 2004) found in Florida rosemary scrub, in recently burned or other disturbed areas (Menges & Kimmich 1996). Fire kills the majority of rosettes and the population recovers through the seedbank (Menges & Kohfeldt 1995). We used demographic data from a single population that forms part of a well-studied meta-population at the Archbold Biological Station, Florida (Appendix A2; Menges & Quintana-Ascencio 2004).

Altering the frequency of fires is one possible management strategy for this endangered species. The recommended fire return interval (FRI) for this species of <15 years (Menges & Quintana-Ascencio 2004) differs from the 15-30 year recommendations for its Florida scrubland habitat (Menges 2007). Alternative management strategies may therefore be required for *Eryngium*. We use perturbation analyses to determine how altering FRIs and the effect of fire on the vital rates affects population growth.

The *Eryngium* IPM (Fig. 5a; Appendix A2) was structured by the natural logarithm of rosette diameter (Menges & Quintana-Ascencio 2004). We assume density independent dynamics to investigate the persistence of the population (Menges & Quintana-Ascencio 2004, see Appendix A5 for model with density dependent recruitment). The intercepts of four vital rates were assumed to be temporally variable (Fig. 5b): survival, growth, flowering probability, and fecundity (the number of flowering stems; Appendix A2). As the demography of *Eryngium* is strongly affected by fire, we modelled the mean of the latent parameter (*Q*) as a linear function of time since fire (*TSF*; Figs 5b & S4). Flowering and fecundity were highly correlated, so the flowering (*ε^f^*) and fecundity (*ε^fe^*) year effects were sampled from a bivariate normal distribution. Sampling these parameters from univariate distributions results in the latent variable failing to fully account for the covariation among the vital rates (Fig. S5). Posterior samples were again drawn using MCMC sampling in JAGS, using runjags (Denwood in review). Weakly informative priors were used (Appendix A2; see Appendix A3 for a comparison with more informative priors). The vital rates were negatively related with TSF, with survival particularly strongly affected (Figs 5b & S6).

**Figure 5:**
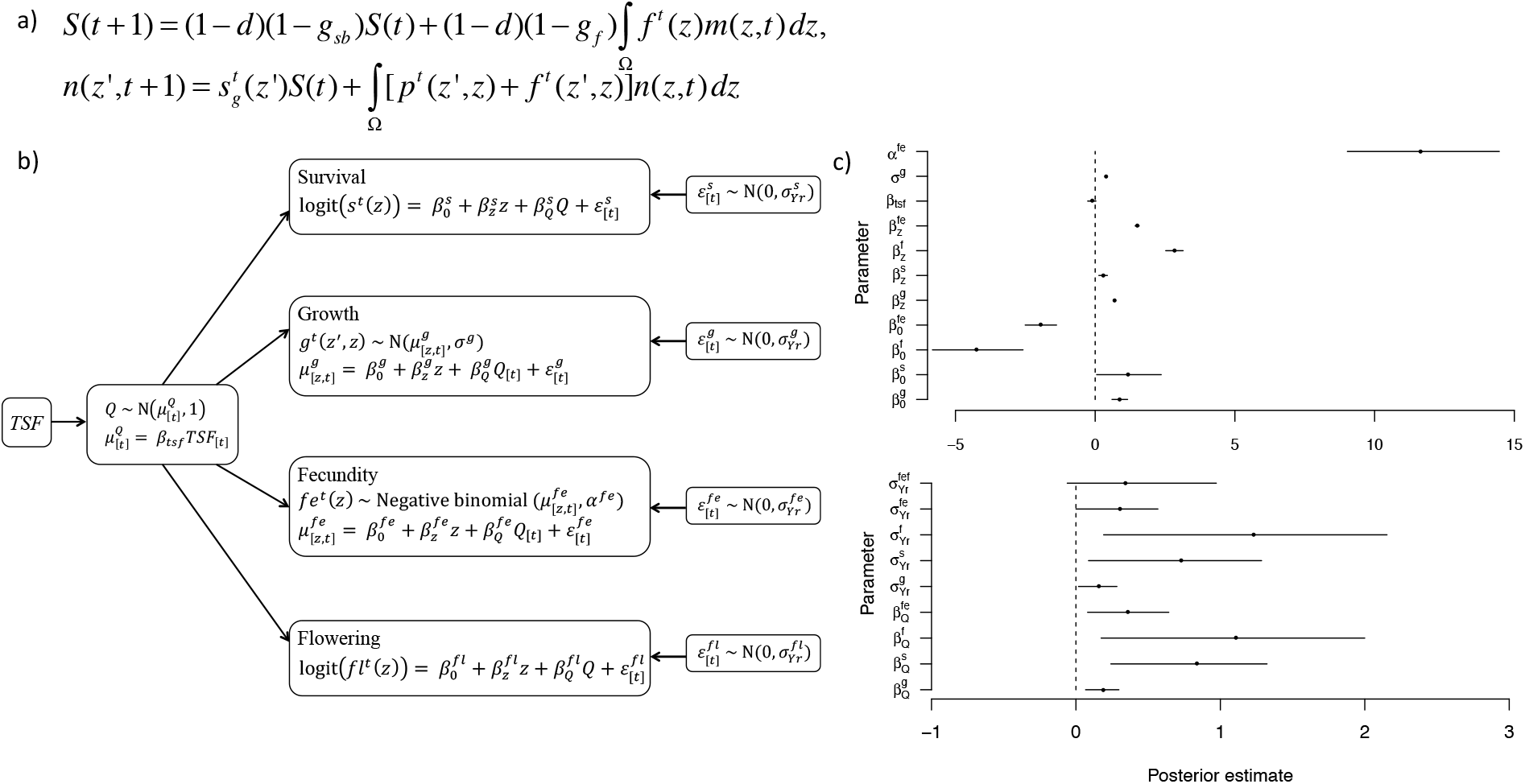
a) *Eryngium* IPM kernel (see Appendix A2). The first term of the seedbank (*S*) equation is those seeds that remain in the seedbank from *t* to *t* + 1; they do not die, with probability 1 - *d*, and do not germinate, with probability 1 -*g_sb_*. The second term refers to seeds produced that year that enter the seedbank. The size distribution of rosettes at *t* + 1 is given by those seeds germinating from the seedbank 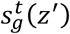, a survival-growth function, *p^t^(z',z)*, of rosettes at *t* and fecundity function, *f^t^(z',z)*. b) Structure of *Eryngium* stochastic vital rates model, including equations for each submodel and b) posteriors for this model for (i) fixed effects and (ii) year effects. *β*_0_ parameters are intercepts, *β_z_* and *β_Q_* parameters are slopes with respect to size (*z*) and the latent parmeter (*Q*) respectively and *ε* are submodel specific year effects. *σ^g^* and *σ^g^* are the standard deviation for the growth and recruitment functions respectively and *α^fe^* is the dispersion parameter for the fecundity function. *σ^Yr^* parameters are the standard deviations of the submodel specific year effects; 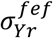 is the covariance between the fecundity and flowering year effects. In b) the points show the means and horizontal lines show the 95% credible intervals for each parameter; the vertical dashed line is at 0.

The posterior means were used to parameterise an IPM. At each iteration the latent parameter () was randomly sampled from a normal distribution with mean *β_tsf_×TSF* and standard deviation of one. Sub-model specific year effects were drawn from normal distributions (bivariate normal for flowering and fecundity), with means of zero and the estimated (co)variances. Estimates of germination probability range from 0 to 0.1 and 0.005 to 0.04 for first (*g_f_*) and second year germination (*g_sb_*) respectively (Quintana-Ascencio & Menges 2000; Menges & Quintana-Ascencio 2004). To select a fertility scenario for the perturbation analyses predicted dynamics using a range of these estimates and of seed mortality probabilities (0.5, 0.7, 0.9) were compared to those observed in the field (Appendix A2). A model with low first year germination (0.0), high germination from the seedbank (0.04) and low seed mortality (0.5) was selected as it was consistent with observed changes in aboveground population growth (Fig. 6a). That is, aboveground populations were predicted to increase immediately following a fire, but not beyond ten years postfire (Menges & Quintana-Ascencio 2004).

**Figure 6:**
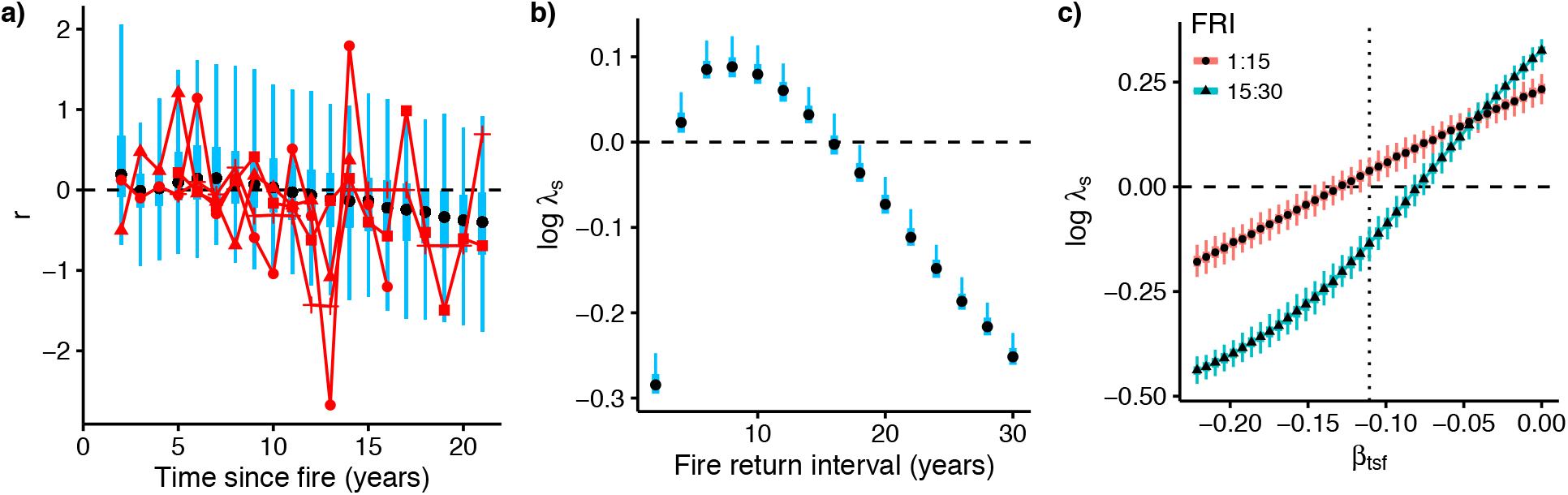
(a) Aboveground population growth rate 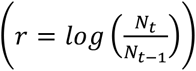 estimated from 1000 simulations of 22 years each for *Eryngium*. Red points are the observed growth rates and mean sizes for the study population and three other populations (site numbers 45, 57 and 91) with similar FRIs. (b) Log *λ_s_* under different burning regimes, (c) effect of changing value of *β_tsf_* on log *λ_s_* under two different FRIs. In all four plots *g_f_ = 0.0, g_sb_ = 0.04* and *d* = 0.5. Points show the median, thicker bars and thinner bars show interquartile range and 95% quantiles respectively. Dotted vertical line in (c) shows the estimated value of *β_tsf_*.

#### Perturbation analyses

The effects of different fire regimes were explored using a range of constant fire return intervals (FRIs) from two to 30 years. Stochastic population growth rates were estimated by iterating 100 populations for 1,000 years; the first 200 years were excluded as transient dynamics. We found populations were likely to decline where the time between fires was too short (c.<4 years), because plants do not produce enough seeds to replenish the seedbank, or too long (c.>15 years; Fig. 6b), as they are outcompeted. This is in accordance with a previous study, using a matrix selection approach, which found an optimal FRI of less than 15 years (Menges & Quintana-Ascencio 2004).

To determine how altering the effect of fire on the vital rates affected population growth the *β_tsf_* parameter was perturbed. This is a measure of how quickly the environment decays as TSF increases; more negative values of this parameter indicate the quality of the environment decreases more quickly following a fire. Stochastic population growth rates were estimated as before, but the fire regimes were varied randomly throughout the simulations (with the same chance of each FRI occurring), either between 1 and 15 years (optimum for *Eryngium*) or between 15 and 30 years (optimum for Florida scrub habitat). Decreasing *β_tsf_* by around 1/3 could make a 15:30 year FRI strategy sustainable for *Eryngium* (Fig. 6c). The effect of altering the temporal decay of the environment is much higher when the FRI is higher, to the extent that decreasing *β_tsf_* sufficiently can make longer FRIs preferable for *Eryngium* (Fig. 6c).

## Discussion

Identifying the environmental drivers of variation in demographic performance is challenging. A variety of approaches have been proposed (e.g. Teller *et al*. 2016; Van der Pol *et al*. 2016), but the performance of any method is limited by the degree of temporal replication available. The mean length for a demographic data set is six years in plants and eleven in animals (Salguero-Gomez *et al*. 2015; Salguero-Gomez *et al*. 2016). Yet, simulations suggest 20-25 years of data are needed to identify environmental drivers, determine the temporal window over which they act and reliably estimate the magnitude of their effects (Teller *et al*. 2016; Van der Pol *et al*. 2016). Efforts to identify drivers in many of these populations will not succeed, forcing population ecologists to assess the likely effects of environmental change using indirect methods. The observation that, in natural populations, different components of demographic performance covary, often positively, (Nur & Sydeman 1999; Jongejans *et al*. 2010; Rotella *et al*. 2012) implies different demographic processes respond (at least in part) to the same drivers. We have demonstrated how a factor analytic (FA) framework can be used to incorporate a temporal axis of environmental variation into a demographic model. The resulting multi-process model—coupled via a latent ‘environmental quality’ variable—requires fewer parameters than its unstructured (UCM) counterpart. In principle, FA models may yield more precise estimates of population growth, though this comes at a potential cost of increased bias when the model is insufficiently flexible. In practice, in our simulation study the UCM and FA approaches yielded comparable estimates of population growth rate. Thus, the main advantage of the FA approach is that it identifies the main axes of demographic variation, which provide a basis for understanding how populations may respond to environmental change.

When it is not possible to explicitly identify environmental drivers of demographic variation, local perturbation analysis of model parameters can be used to explore the potential response of a population to environmental change (Rees & Ellner 2009). These analyses typically consider each parameter in turn, assessing its effect on metrics such as population growth rate while holding all else constant. However, the existence of (positive) temporal correlations among demographic processes suggests multiple processes respond in a coordinated manner to environmental change. An FA model allows us to identify the potential axis of change and, by focusing perturbation analyses on this axis, makes population level predictions under different environmental conditions possible. For example, this allowed us to make predictions on how life histories may evolve under putative environmental conditions. We identified how the environmental quality would have to change for *Carduus* to alter its flowering strategy; showing that increases in its average vital rates, in particular survival, will lead to selection for a perennial life history. Whilst we focus here on temporal variation this approach could also be used to predict joint demographic responses to spatial variation (Elderd & Miller 2016).

The key limitation is that this interpretation of the FA model assumes temporal covariances are largely environmentally driven. This may not be true if individuals substantially adjust their allocation strategy in response to environmental conditions. Negative correlations among the vital rates may exist due to life history trade-offs between vital rates, where for example resources are invested in survival or reproduction to the detriment of growth (Koenig & Knops 1998). Negative correlations appear relatively rare however (Jongejans *et al*. 2010), and where they do exist are sometimes attributable to opposing responses to environmental drivers (e.g. Knops, Koenig & Carmen 2007). This suggests the magnitude of trade-off effects is generally small compared to that of environmental effects, though life history tradeoffs may still attenuate environmental driver(s) of covariation.

Explicitly incorporating environmental drivers allows population responses to management strategies or anticipated environmental change to be predicted (e.g. Gotelli & Ellison 2006; Isaza *et al*. 2016). The FA approach can simplify the process of incorporating such drivers, as they can be included into a single model of the shared environmental axis. Where explicit environmental drivers (e.g. population density or temperature) can be identified, these are typically considered on a process-specific basis, by constructing separate models for survival, reproduction, growth, and recruitment (e.g. Dahlgren, Ostergard & Ehrlen 2014; Williams *et al*. 2015). This would require the addition of four time-since-fire slope parameters in our *Eryngium* case study, one for each temporally variable vital rate (e.g. Evans, Holsinger & Menges 2010). Instead we introduced a higher-level model, decomposing the shared axis of environmental variation into explained and residual components of variation. Thus the effects of time since fire on all four vital rates were incorporated with the addition of a single parameter. This allowed us to evaluate the population level effects of two alternative management strategies; that is altering the disturbance regime or ameliorating the environment to decrease the rate of decay in environmental quality following a disturbance. We found that whilst the optimum FRI for *Eryngium* is less than 15 years, decreasing the rate of environmental decay could lead to persistent populations under 15-30 year fire regimes (the recommended FRI for the Florida scrub habitat; Menges 2007).

Similar approaches to our analysis have been used previously (Evans, Holsinger & Menges 2010; Evans & Holsinger 2012; Elderd & Miller 2016). However, in previous models the slope terms for the environmental quality parameter (*β_Q_*) were constrained to a value of +1 or -1 among a set of demographic models. These usually operate on different scales, for example probabilities, such as survival and flowering are typically estimated on a logit scale, whereas fecundity is generally estimated on a log scale; a unit change on these two scales cannot be meaningfully compared. Moreover, differences in the magnitude of the effect of temporal variation among the vital rates were accounted for by the process specific year effects (*ε*). Thus the main advantage of the FA approach is lost, as the latent variable cannot be conceived as a simple measure of overall environmental quality.

Adopting a Bayesian approach has a number of benefits (Elderd & Miller 2016), for example allowing the effects of difference sources of uncertainty to be quantified (Evans, Holsinger & Menges 2010). Uncertainty is likely to be very high for most data sets (Metcalf *et al*. 2015). Parameter uncertainty can have important ecological implications, for example failing to account for it may underestimate the risk of extinction (Ludwig 1996). Using a Bayesian approach also allows for posterior predictive checks (Gelman *et al*. 2004). These are particularly important when fitting very constrained models, for example when assuming the temporal covariation in the vital rates may be explained by a single environmental axis. Sometimes, as in the *Eryngium* case study, additional axes may be necessary to fully account for the covariation in the vital rates. We recommend starting with a simple model structure and slowly adding in complexity.

Rapid levels of environmental change have increased interest in determining how population processes respond to environmental stochasticity (Stenseth *et al*. 2002; Evans 2012). However, the longterm individual level data needed to accurately quantify such responses are often lacking, especially for rare species. Where positive covariances exist among vital rates these can be exploited under a FA approach to allow predictions on the joint responses of vital rates to environmental variation. Where insufficient data exist to identify environmental drivers the FA approach may offer the best alternative for predicting population responses to environmental change.

## Acknowledgements

BJH was funded by a NERC and University of Sheffield PhD studentship. DZC was funded by a NERC fellowship (NE/I022027/1). *Eryngium* data were provided by the Archbold Biological Station.

## Authors’ contributions

AWS, PFQ-A, and ESM collected the data; analysis was carried out by BJH with guidance from DZC and MR; BJH and DZC led the writing of the manuscript. All authors contributed critically to the drafts and gave final approval.

## References

Benton, T.G. & Grant, A. (1996) How to keep fit in the real world: Elasticity analyses and selection pressures on life histories ln a variable environment. American Naturalist, 147, 115–139.

Boyce, M.S., Haridas, C.V., Lee, C.T. & Demography, N.S. (2006) Demography in an increasingly variable world. Trends in Ecology & Evolution, 21, 141–148.

Caswell, H. (2001) Matrix population models: Construction, analysis, and interpretation. Matrix population models: Construction, analysis, and interpretation, i-xxii, 1–722.

Childs, D.Z., Metcalf, C.J.E. & Rees, M. (2010) Evolutionary bet-hedging in the real world: empirical evidence and challenges revealed by plants. Proceedings of the Royal Society B-Biological Sciences, 277, 3055–3064.

Childs, D.Z., Rees, M., Rose, K.E., Grubb, P.J. & Ellner, S.P. (2003) Evolution of complex flowering strategies: an age- and size-structured integral projection model. Proceedings of the Royal Society B-Biological Sciences, 270, 1829–1838.

Childs, D.Z., Rees, M., Rose, K.E., Grubb, P.J. & Ellner, S.P. (2004) Evolution of size-dependent flowering in a variable environment: construction and analysis of a stochastic integral projection model. Proceedings of the Royal Society B-Biological Sciences, 271, 425–434.

Clutton-Brock, T.H. & Pemberton, J.M. (2004) Soay Sheep: Dynamics and Selection in an Island Population.

Cohen, D. (1966) Optimizing reproduction in a randomly varying environment. Journal of Theoretical Biology, 12, 119-&.

Coulson, T. (2012) Integral projections models, their construction and use in posing hypotheses in ecology. Oikos, 121, 1337–1350.

Crone, E.E., Ellis, M.M., Morris, W.F., Stanley, A., Bell, T., Bierzychudek, P., … & Menges, E.S. (2013) Ability of Matrix Models to Explain the Past and Predict the Future of Plant Populations. Conservation Biology, 27, 968–978.

Dahlgren, J.P., Ostergard, H. & Ehrlen, J. (2014) Local environment and density-dependent feedbacks determine population growth in a forest herb. Oecologia, 176, 1023–1032.

Darling, E.S. & Cote, I.M. (2008) Quantifying the evidence for ecological synergies. Ecology Letters, 11, 1278–1286.

Denwood, M.J. (in review) runjags: An R package providing interface utilities, parallel computing methods and additional distributions for MCMC models in JAGS. Journal of Statistical Software.

Doak, D.F., Morris, W.F., Pfister, C., Kendall, B.E. & Bruna, E.M. (2005) Correctly estimating how environmental stochasticity influences fitness and population growth. American Naturalist, 166, E14–E21.

Ehrlen, J., Morris, W.F., von Euler, T. & Dahlgren, J.P. (2016) Advancing environmentally explicit structured population models of plants. Journal of Ecology, 104, 292–305.

Elderd, B.D. & Miller, T.E.X. (2016) Quantifying demographic uncertainty: Bayesian methods for integral projection models. Ecological Monographs, 86, 125–144.

Ellner, S.P., Childs, D.Z. & Rees, M. (2016) Data-driven modelling of structured populations: A practical guide to the Integral Projection Model. Springer, Switzerland.

Evans, M.E.K. & Holsinger, K.E. (2012) Estimating covariation between vital rates: A simulation study of connected vs. separate generalized linear mixed models (GLMMs). Theoretical Population Biology, 82, 299–306.

Evans, M.E.K., Holsinger, K.E. & Menges, E.S. (2010) Fire, vital rates, and population viability: a hierarchical Bayesian analysis of the endangered Florida scrub mint. Ecological Monographs, 80, 627–649.

Evans, M.R. (2012) Modelling ecological systems in a changing world. Philosophical Transactions of the Royal Society B-Biological Sciences, 367, 181–190.

Fieberg, J. & Ellner, S.P. (2001) Stochastic matrix models for conservation and management: a comparative review of methods. Ecology Letters, 4, 244–266.

Gelman, A., Carlin, J.B., Stern, H.S. & Rubin, D.B. (2004) Bayesian Data Analysis, Second Edition. Chapman and Hall/CRC.

Gotelli, N.J. & Ellison, A.M. (2006) Forecasting extinction risk with nonstationary matrix models. Ecological Applications, 16, 51–61.

Grace, J.B., Anderson, T.M., Olff, H. & Scheiner, S.M. (2010) On the specification of structural equation models for ecological systems. Ecological Monographs, 80, 67–87.

Grace, J.B. & Bollen, K.A. (2008) Representing general theoretical concepts in structural equation models: the role of composite variables. Environmental and Ecological Statistics, 15, 191–213.

Inchausti, P. & Weimerskirch, H. (2001) Risks of decline and extinction of the endangered Amsterdam albatross and the projected impact of long-line fisheries. Biological Conservation, 100, 377–386.

Isaza, C., Martorell, C., Cevallos, D., Galeano, G., Valencia, R. & Balslev, H. (2016) Demography of Oenocarpus bataua and implications for sustainable harvest of its fruit in western Amazon. Population Ecology, 58, 463–476.

Jenkins, D., Watson, A. & Miller, G.R. (1963) Population studies on red grouse, Lagopus-lagopus-scoticus (Lath), in northeast Scotland. Journal of Animal Ecology, 32, 317–376.

Jongejans, E. & De Kroon, H. (2005) Space versus time variation in the population dynamics of three co-occurring perennial herbs. Journal of Ecology, 93, 681–692.

Jongejans, E., de Kroon, H., Tuljapurkar, S. & Shea, K. (2010) Plant populations track rather than buffer climate fluctuations. Ecology Letters, 13, 736–743.

Knops, J.M.H., Koenig, W.D. & Carmen, W.J. (2007) Negative correlation does not imply a tradeoff between growth and reproduction in California oaks. Proceedings of the National Academy of Sciences of the United States of America, 104, 16982–16985.

Koenig, W.D. & Knops, J.M.H. (1998) Scale of mast-seeding and tree-ring growth. Nature, 396, 225–226.

Ludwig, D. (1996) Uncertainty and the assessment of extinction probabilities. Ecological Applications, 6, 1067–1076.

Menges, E.S. (2007) Integrating demography and fire management: an example from Florida scrub. Australian Journal of Botany, 55, 261–272.

Menges, E.S. & Kimmich, J. (1996) Microhabitat and time-since-fire: Effects on demography of Eryngium cuneifolium (Apiaceae), a Florida scrub endemic plant. American Journal of Botany, 83, 185–191.

Menges, E.S. & Kohfeldt, N. (1995) Life history strategies of Florida scrub plants in relation to fire. Bulletin of the Torrey Botanical Club, 122, 282–297.

Menges, E.S. & Quintana-Ascencio, P.F. (2004) Population viability with fire in Eryngium cuneifolium: Deciphering a decade of demographic data. Ecological Monographs, 74, 79–99.

Metcalf, C.J.E., Ellner, S.P., Childs, D.Z., Roberto, S.-G., Merow, C., McMahon, S.M., Jongejans, E. & Rees, M. (2015) Statistical modelling of annual variation for inference on stochastic population dynamics using Integral Projection Models. Methods in Ecology and Evolution, 6, 1007–1017.

Morris, W.F. & Doak, D.F. (2002) Quantitative Conservation Biology: Theory and Practice of Population Viability Analysis. Quantitative Conservation Biology: Theory and Practice of Population Viability Analysis, i-xvi, 1–480.

Nur, N. & Sydeman, W.J. (1999) Survival, breeding probability and reproductive success in relation to population dynamics of Brandt’s cormorants Phalacrocorax penicillatus. Bird Study, 46, 92–103.

Ohlberger, J., Scheuerell, M.D. & Schindler, D.E. (2016) Population coherence and environmental impacts across spatial scales: a case study of Chinook salmon. Ecosphere, 7.

Ottersen, G., Planque, B., Belgrano, A., Post, E., Reid, P.C. & Stenseth, N.C. (2001) Ecological effects of the North Atlantic Oscillation. Oecologia, 128, 1–14.

Parmesan, C., Burrows, M.T., Duarte, C.M., Poloczanska, E.S., Richardson, A.J., Schoeman, D.S. & Singer, M.C. (2013) Beyond climate change attribution in conservation and ecological research. Ecology Letters, 16, 58–71.

Plummer, M. (2003) JAGS: a program for analysis of Bayesian graphical models using Gibbs sampling. pp. 125. Proceedings of the 3rd international workshop on distributed statistical computing. Technische Universit at Wien, Wien, Austria.

Popay, A.I. & Medd, R.W. (1990) The biology of Australian weeds 21. Carduus-nutans-ssp-nutans. Plant Protection Quarterly, 5, 3–13.

Post, E. & Stenseth, N.C. (1999) Climatic variability, plant phenology, and northern ungulates. Ecology, 80, 1322–1339.

Qiu, W. & Joe, H. (2015) clusterGeneration: Random Cluster Generation (with Specified Degree of Separation). R package version 1.3.4. http://CRAN.R-project.org/package=clusterGeneration.

Quintana-Ascencio, P.F. & Menges, E.S. (2000) Competitive abilities of three narrowly endemic plant species in experimental neighborhoods along a fire gradient. American Journal of Botany, 87, 690–699.

R Core Team (2016) R: A language and environment for statistical computing. R Foundation for Statistical Computing, Vienna, Austria.

Rees, M., Childs, D.Z., Metcalf, J.C., Rose, K.E., Sheppard, A.W. & Grubb, P.J. (2006) Seed dormancy and delayed flowering in monocarpic plants: Selective interactions in a stochastic environment. American Naturalist, 168, E53–E71.

Rees, M. & Ellner, S.P. (2009) Integral projection models for populations in temporally varying environments. Ecological Monographs, 79, 575–594.

Rotella, J.J., Link, W.A., Chambert, T., Stauffer, G.E. & Garrott, R.A. (2012) Evaluating the demographic buffering hypothesis with vital rates estimated for Weddell seals from 30 years of mark-recapture data. Journal of Animal Ecology, 81, 162–173.

Salguero-Gomez, R., Jones, O.R., Archer, C.R., Bein, C., de Buhr, H., Farack, C., … & Vaupel, J.W. (2016) COMADRE: a global data base of animal demography. Journal of Animal Ecology, 85, 371–384.

Salguero-Gomez, R., Jones, O.R., Archer, C.R., Buckley, Y.M., Che-Castaldo, J., Caswell, H., … & Vaupel, J.W. (2015) The COMPADRE Plant Matrix Database: an open online repository for plant demography. Journal of Ecology, 103, 202–218.

Stenseth, N.C. & Mysterud, A. (2005) Weather packages: finding the right scale and composition of climate in ecology. Journal of Animal Ecology, 74, 1195–1198.

Stenseth, N.C., Mysterud, A., Ottersen, G., Hurrell, J.W., Chan, K.S. & Lima, M. (2002) Ecological effects of climate fluctuations. Science, 297, 1292–1296.

Teller, B.J., Adler, P.B., Edwards, C.B., Hooker, G. & Ellner, S.P. (2016) Linking demography with drivers: climate and competition. Methods in Ecology and Evolution, 7, 171–183.

Thorson, J.T., Scheuerell, M.D., Shelton, A.O., See, K.E., Skaug, H.J. & Kristensen, K. (2015) Spatial factor analysis: a new tool for estimating joint species distributions and correlations in species range. Methods in Ecology and Evolution, 6, 627–637.

Tomimatsu, H. & Ohara, M. (2010) Demographic response of plant populations to habitat fragmentation and temporal environmental variability. Oecologia, 162, 903–911.

Tuljapurkar, S. (1990) Delayed reproduction and fitness in variable environments. Proceedings of the National Academy of Sciences of the United States of America, 87, 1139–1143.

Van der Pol, M., Bailey, L.D., McLean, N., Rijsdijk, L., Lawson, C.R. & Brouwer, L. (2016) Identifying the best climatic predictors in ecology and evolution. Methods in Ecology and Evolution, 7, 1246–1257.

Vindenes, Y., Edeline, E., Ohlberger, J., Langangen, O., Winfield, I.J., Stenseth, N.C. & Vollestad, L.A. (2014) Effects of Climate Change on Trait-Based Dynamics of a Top Predator in Freshwater Ecosystems. American Naturalist, 183, 243–256.

Wardle, D.A., Nicholson, K.S. & Rahman, A. (1992) Influence of pasture grass and legume swards on seedling emergence and growth of Carduus-nutans and Cirsium-vulgare. Weed Research, 32, 119–128.

Williams, J.L., Jacquemyn, H., Ochocki, B.M., Brys, R. & Miller, T.E.X. (2015) Lifehistory evolution under climate change and its influence on the population dynamics of a long-lived plant. Journal of Ecology, 103, 798–808.

Zuur, A.F., Fryer, R.J., Jolliffe, I.T., Dekker, R. & Beukema, J.J. (2003) Estimating common trends in multivariate time series using dynamic factor analysis. Environmetrics, 14, 665–685.

